# Dynamic shifts of visual and saccadic signals in prefrontal cortical regions 8Ar and FEF

**DOI:** 10.1101/817478

**Authors:** Sanjeev B. Khanna, Jonathan A. Scott, Matthew A. Smith

**Affiliations:** Dept. of Ophthalmology, Univ. of Pittsburgh, Pittsburgh, PA, USA; Center for the Neural Basis of Cognition, Univ. of Pittsburgh, Pittsburgh, PA, USA; Dept. of Bioengineering, Univ. of Pittsburgh, Pittsburgh, PA, USA; Dept. of Biomedical Engineering, Carnegie Mellon University, Pittsburgh, PA, USA; Carnegie Mellon Neuroscience Institute, Pittsburgh, PA, USA

**Keywords:** prefrontal cortex, saccade, working memory, receptive field, dynamics

## Abstract

Active vision is a fundamental process by which primates gather information about the external world. Multiple brain regions have been studied in the context of simple active vision tasks in which a visual target’s appearance is temporally separated from saccade execution. Most neurons have tight spatial registration between visual and saccadic signals, and in areas such as prefrontal cortex (PFC) some neurons show persistent delay activity that links visual and motor epochs and has been proposed as a basis for spatial working memory. Many PFC neurons also show rich dynamics, which have been attributed to alternative working memory codes and the representation of other task variables. Our study investigated the transition between processing a visual stimulus and generating an eye movement in populations of PFC neurons in macaque monkeys performing a memory guided saccade task. We found that neurons in two subregions of PFC, the frontal eye fields (FEF) and area 8Ar, differed in their dynamics and spatial response profiles. These dynamics could be attributed largely to shifts in the spatial profile of visual and motor responses in individual neurons. This led to visual and motor codes for particular spatial locations that were instantiated by different mixtures of neurons, which could be important in PFC’s flexible role in multiple sensory, cognitive, and motor tasks.

**New and Noteworthy:** A central question in neuroscience is how the brain transitions from sensory representations to motor outputs. The prefrontal cortex contains neurons that have long been implicated as important in this transition and in working memory. We found evidence for rich and diverse tuning in these neurons, that was often spatially misaligned between visual and saccadic responses. This feature may play an important role in flexible working memory capabilities.

## Introduction

The process of gathering information about the external world and acting on it necessitates sensorimotor integration and is crucial for survival and adaptation to the environment. In a simple idealized organism, visual space could be directly mapped onto motor responses. This type of architecture would be well suited to direct orienting responses in a hardwired fashion, where a stimulus and resulting movement need to be processed and generated rapidly. Within the oculomotor domain, one example of a behavior that could be accomplished by such a direct mapping is the generation of a rapid, ballistic eye movement (saccade) to a visual stimulus. Neurons in the superior colliculus (SC), a midbrain region tightly coupled to the brain stem circuits that move the eyes, have overlapping visual and movement-related activity (Wurtz & Goldberg 1972). Such a tight alignment might be ideal for the rapid translation of perception to action, and is particularly important in cognitive functions such as attention that involve a tight interaction between visual and saccadic maps (Krauzlis et al 2013). However, in many instances, stimuli must be represented first and acted on later with one of several motor output modalities in the context of different cognitive constraints. The means by which such flexible sensorimotor behavior is achieved is an important mystery in neuroscience.

Sensorimotor signals have been extensively studied in the context of eye movements, which are a critical part of primate behavior but also have the advantage of high repeatability and a limited number of degrees of freedom. In particular, tasks in which the onset of the visual stimulus and the eye movement are temporally separated (such as the memory-guided saccade, or MGS) have been used to study the transformation from perception to action. Neurons in oculomotor regions of the cortex typically do not respond exclusively to a visual stimulus or an eye movement. Instead, they demonstrate a wide variety of activity profiles relating to their visuomotor response properties and the timing and duration of their activity. Two prefrontal cortical regions which contain neurons with these diverse response properties are the frontal eye fields (FEF) and the pre-arcuate gyrus (Bullock et al 2017, Kiani et al 2015, Preuss & Goldman-Rakic 1991, Schall et al 1995). In both regions, neurons respond to visual stimuli, eye movements, or both in varying degrees (Boch & Goldberg 1989, Bruce & Goldberg 1985, Funahashi et al 1991). Additionally, some neurons fire transient bursts often aligned to stimulus or saccade onset, while others maintain their activity throughout the period between the visual stimulus and saccade (Funahashi et al 1989, Funahashi et al 1990, Fuster & Alexander 1971). Activity that is elevated and sustained during the entire delay epoch (the period of time after the spatial location is stored in memory but before it must be retrieved), referred to as persistent activity, has been proposed to underlie spatial working memory (Goldman-Rakic 1995). However, while some neurons maintain relatively constant activity in the delay period, many others exhibit changes over time such as ramping up or down or shifts in preference. This has led to ongoing debate about the nature of neural signals related to working memory (for review, see Constantinidis et al (2018) and Lundqvist et al (2018)) and also provides insight into the transition between visual and motor signals.

The predominant observation of FEF neurons has been of alignment between sensory and motor responses in representing the contralateral visual field (Bruce & Goldberg 1985), similar to SC. However, a small subset of FEF neurons have ipsilateral receptive fields (Crapse & Sommer 2009, Schall 1991) and can even show misalignment between their delay period and peri-saccadic tuning (Lawrence et al 2005). In 8Ar, there are also neurons with ipsilateral receptive fields (Mikami et al 1982, Suzuki & Azuma 1983) and bilateral responses (Bullock et al 2017). Some PFC neurons change their tuning during the delay epoch (Parthasarathy et al 2017, Spaak et al 2017), which has led some to propose alternatives to the persistent activity model of working memory, such as “activity silent” mechanisms (Stokes 2015), or oscillatory dynamics (Lundqvist et al 2016). Setting aside the question of how working memory is stored, there is abundant evidence that neurons in PFC represent a myriad of perceptual and task-related variables, such as reward (Leon & Shadlen 1999, Watanabe 1996), abstract rules (Wallis et al 2001), time during the delay (Jun et al 2010, Spaak et al 2017), previous trial outcome (Donahue & Lee 2015), and stimulus shape and color (Riley et al 2017). A misalignment of visual and eye movement signals could be the consequence of, and potentially beneficial for, a flexible coding architecture in which multiple perceptual inputs (e.g., visual or auditory) are associated with multiple motor outputs (e.g., a saccade or a reach). In this case, the dynamics of activity during the delay period may not solely represent the evolution of a working memory signal, but also the transition between two different representations in the neuronal population. We wondered if and how the visual and saccadic signals align in FEF and 8Ar, and whether there were systematic rules by which neurons shifted their response profiles.

To answer these questions, we recorded from groups of 8Ar and FEF neurons in macaque monkeys performing a memory guided saccade task. The reliable timing of the task and saccadic response allowed us to isolate visual and motor signals. We first characterized the receptive field structure of 8Ar neurons in response to a briefly flashed visual stimulus as well as the motor response field around the time of the saccade. Importantly, we used a dense mapping of space to achieve a high-resolution spatial response profile. We found a remarkable amount of diversity, both spatially and temporally, in the response properties of 8Ar neurons. A key pattern in this diversity was spatial and temporal opponency – many neurons were suppressed at spatial locations opposite their preferred response, and their preferences shifted over time from shortly after stimulus onset to the time of the saccade, sometimes to the opposite hemifield. To quantitatively assess at the population level the observations that we made in individual neurons, we measured the ability to decode the target location in 8Ar and FEF neurons. We found that the diverse response properties observed in 8Ar are less prominent in FEF, and that the population code in 8Ar is more dynamic, meaning that the visual and motor codes were less aligned. Taken together, our results are consistent with a neural circuit structure in which the visual and saccadic representations are gradually segregated with increasing distance from the motor output. This may aid in 8Ar playing a flexible role in various types of sensorimotor behavior and cognitive states, associating a variety of sensory inputs with potential motor outputs.

### Materials and Methods

### Neuronal recordings

#### Surgical Preparation

A 96-electrode “Utah” Array (Blackrock Microsystems, Salt Lake City, UT) was implanted into two adult, male rhesus macaques (*Macaca mulatta*) in dorsolateral prefrontal cortex using sterile surgical techniques under isoflurane anesthesia. The array was implanted in right 8Ar for Monkey Pe and left 8Ar for Monkey Wa, on the pre-arcuate gyrus immediately anterior to the arcuate sulcus and medial to the principal sulcus (Figure 1A). For FEF recordings, two adult male rhesus macaques (*Macaca mulatta*; monkeys Ro and Wi) were surgically implanted with FEF recording chambers (aimed for the anterior bank of the arcuate sulcus, centered at stereotaxic coordinates: 25 anterior, 20 lateral) and extracellular activity was recorded with a 16-electrode linear microelectrode array (U-Probe, Plexon, Dallas TX) with contacts spaced 150 µm apart. Linear arrays were lowered into FEF daily using a custom designed mechanical microdrive (Laboratory for Sensorimotor Research, National Eye Institute, Bethesda, MD) through a plastic grid with 1 mm spacing. The FEF recordings included in this analysis were part of a larger dataset previously published (Khanna et al 2019). The location of FEF was first identified by physiological response properties to visual stimuli and saccades, and then confirmed through microstimulation. Recording sites were considered to be in FEF if saccades could be reliably (>50%) evoked using low threshold microstimulation (≤ 50 µA, 0.25 ms pulse width, 70 ms pulse train duration, 350 Hz stimulation frequency) (Bruce et al 1985) at that location or at an immediately neighboring grid location (1mm away). Of note, we were not able to induce eye movements by stimulating any electrodes on the 8Ar arrays, even with microstimulation up to currents of 150 µA. The head was immobilized during recordings with a titanium headpost attached to the skull with titanium screws, implanted in a separate procedure before the array or chamber implants. All procedures were approved by the Institutional Animal Care and Use Committee of the University of Pittsburgh and complied with guidelines set forth in the National Institute of Health’s *Guide for the Care and Use of Laboratory Animals*.

**Figure 1:**
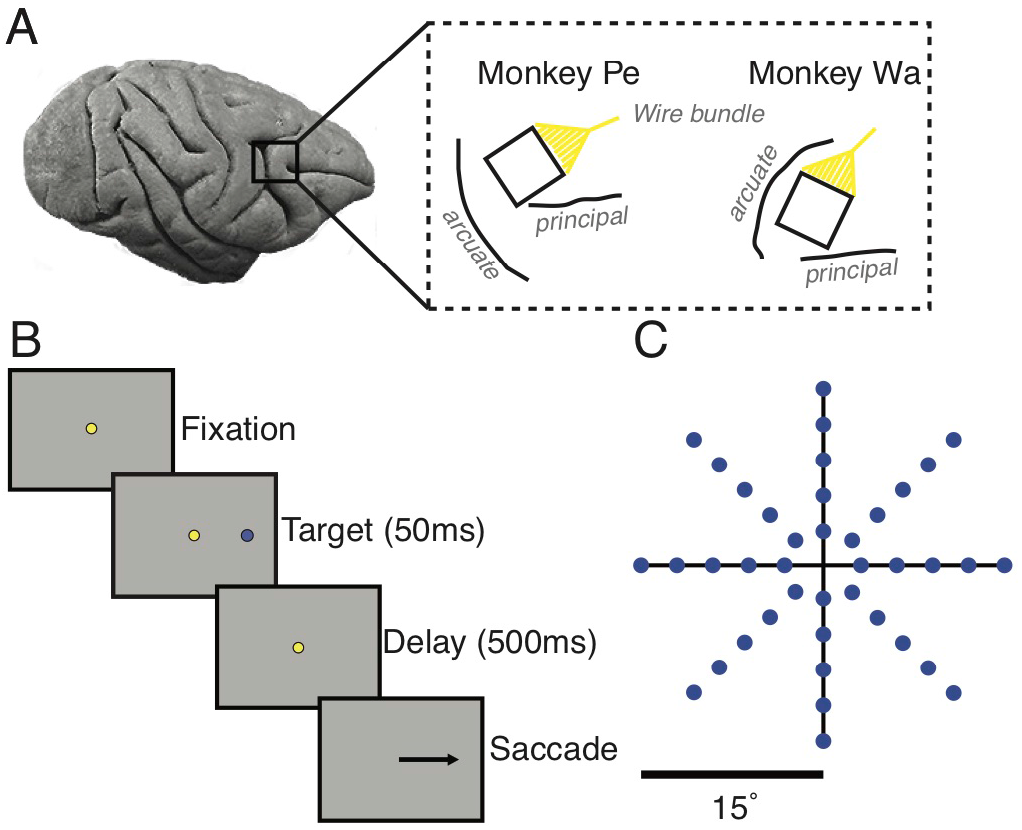
Electrode array locations and task. A) 96 channel Utah arrays were placed in dorsolateral prefrontal cortex (8Ar) on the prearcuate gyrus, anterior to the arcuate sulcus and medial to the principal sulcus. The line drawings indicate visible sulcal patterns through the durotomy and are not meant to represent the full extent of the arcuate and principal sulci. B) Memory guided saccade task. Each trial began with the subject fixating on a central dot. After 200 ms of fixation, a target appeared briefly in the periphery for 50 ms. Following a delay of 500 ms, the fixation point was extinguished, signaling the subject to saccade to the remembered location of the target. C) Targets appeared at 1 of 40 locations, varying in amplitude and direction (only 8 directions with a single amplitude were used for FEF).

#### Data Collection

Stimuli were displayed on a 21” cathode ray tube monitor with a resolution of 1024×768 pixels and a refresh rate of 100 Hz at viewing distance of 36 cm. Stimuli were generated using custom software written in MATLAB (MathWorks, Natick, MA) with the Psychophysics Toolbox extensions (Brainard 1997, Kleiner et al 2007, Pelli 1997). Eye position was tracked monocularly using an infrared system at 1000 Hz resolution (EyeLink 1000, SR Research, Mississauga, Canada). In both the FEF and 8Ar recordings, extracellular activity was recorded from the array, band-pass filtered (0.3 – 7,500 Hz), digitized at 30 kHz, and amplified by a Grapevine system (Ripple, Salt Lake City, UT). Waveforms that exceeded a threshold were saved and stored for offline wave classification. The threshold was set by taking a value (typically −3) and multiplying it by the root mean squared noise measured on each channel. Waveforms were automatically sorted using a competitive mixture decomposition algorithm (Shoham et al 2003) and later refined manually based on waveform shape characteristics and inter-spike interval distributions using custom time amplitude window discrimination software written in MATLAB (https://github.com/smithlabvision/spikesort).

After the waveforms were sorted, the signal-to-noise ratio (SNR) was calculated for each identified unit as the ratio of the average waveform amplitude to the standard deviation of the waveform noise (Kelly et al 2007). We considered only candidate units with an SNR above 2.5 as isolated single neurons for the purpose of further analysis. This resulted in a total of 2511 neurons across 39 recording sessions in 8Ar (Monkey Wa: 1179 units, 20 sessions; Monkey Pe: 1332 units, 19 sessions) and 889 neurons across 50 sessions in FEF (Monkey Wi: 305 units, 14 sessions; Monkey Ro: 584 units, 36 sessions). We did not attempt to determine whether the same units were recorded across multiple days with the Utah array recordings. It is likely that this did occur in some cases, although our recording sessions from 8Ar were often spaced out by a week or more as the animals were also performing an unrelated experiment at the same time. Results from a single session from each animal are shown in Figure 11, and show the representative features of the data from the full population analysis. In FEF, because the U-Probe was inserted in each recording session and in multiple different chamber locations, recording from the same unit across days was not a concern.

#### Experimental design and statistical analysis

##### Behavioral task

Monkeys performed a standard memory guided saccade (MGS) task (Figure 1B) (Hikosaka & Wurtz 1983). The trial commenced when the subject fixated a small blue dot (0.5° diameter) at the center of the screen. For 8Ar recordings, after fixation was established (200 ms), a target appeared in the periphery at one of eight angular directions (0°, 45°, 90°, 135°, 180°, 225°, 270°, 315°) and one of five eccentricities (5°, 7.5°, 9.9°, 12.3°, 14.7°) (40 total possible locations, Figure 1C) for 50 ms. The animal was required to maintain fixation for 500 ms after the target was extinguished, at which point the central fixation point would disappear, signaling the animal to saccade to the remembered location of the stimulus. The monkey had 500 ms to initiate the saccade, and once it had been initiated (defined as the monkey’s eye position leaving a window 1.8° in diameter around the fixation point) the monkey’s eye position had to reach the saccade target within 200 ms and maintain gaze within 2.7° of the location for 150 ms to receive a liquid reward. Each block consisted of pseudorandomized presentations of all 40 conditions, with at least 40 blocks gathered per session (average 58). For 4 of the 20 sessions in Monkey Wa, the angular directions and target eccentricities were different (angles 26°, 71°, 116°, 161°, 206°, 251°, 296°, 341°; amplitudes 2.6°, 3.9°, 5.2°, 6.5°, 7.8°). Data from these sessions were included in population analyses when possible by using the large-amplitude trials (7.8°) but were not included in the population average response field analyses (Figure 4A, B). For FEF recordings, the same behavioral task (MGS) was used, with stimuli appearing at one of eight angular directions (0°, 45°, 90°, 135°, 180°, 225°, 270°, 315°) but at only one amplitude (10°). The fixation time before target onset (200 ms) and target duration (50 ms) were equal to 8Ar, however the delay epoch was 600 ms (as opposed to 500ms for 8Ar). Each block consisted of pseudorandomized presentations of all eight conditions, with at least 50 blocks gathered per session (average 132). On a subset of days, after the fixation point was extinguished and the monkey began its saccade, the target was re-illuminated to aid in saccade completion. The analyses presented here were not affected by this target because all analysis windows were constructed to end prior to any possible visual transient in response to this target.

##### Neuron selection

All neural firing rates were measured during stimulus presentation, the memory epoch, and the perisaccadic epoch. To determine the ideal response epoch in which to measure tuning, we used a method described in Smith et al (2005) in which the variance (across the 40 conditions) was calculated for each neuron in a sliding window of 50 ms. For a neuron tuned to the spatial location of the visual stimulus or the saccade, the variance is largest when the window is aligned to the latency of the neuron (when it exhibits that tuning in the form of spatially variable responses). To isolate the visual and perisaccadic responses, we measured the latency of the visual response from 0 ms to 400 ms after stimulus onset for all neurons. These outer boundary values were determined by visually examining the PSTHs of individual neurons and the population response. Similarly, the saccade response was measured 100 ms before to 50 ms after the saccade. Once the optimal window was identified for each neuron, we determined whether the neuron had significant (p < .01) spatial tuning in that response window using a Kruskal-Wallis one-way analysis of variance on the average firing rates with location as the factor.

##### Response field calculation

The center of each neuron’s response field during the visual and saccade epoch was calculated as follows. For each stimulus location, activity during the time epoch desired (visual, saccade, or across the delay in a sliding window analysis) was baseline-corrected by subtracting the average activity across all conditions (since they were the same prior to stimulus onset) from 30 ms to 180 ms after fixation was established (170 ms to 20 ms before stimulus onset). The resulting baseline subtracted activity was averaged across the trial srepeats for each condition, and then linearly interpolated to obtain a map with a resolution of 0.25° x 0.25°. This map was smoothed using a gaussian filter with a standard deviation of 1°. The center of the response field was defined as the center of mass for all locations with responses ≥75% of the maximum for a given response field map (Zirnsak et al 2014). Only responses above baseline (as opposed to responses suppressed below baseline) were considered for the center of mass calculation. All response field spatial maps are displayed such that the left side of each image corresponded to the contralateral visual hemifield. This required reflecting the spatial response maps for Monkey Wa (where recordings were made from an array implanted in left 8Ar), so that data from both monkeys were displayed with the same coordinate frame.

##### Visual/motor index calculation

To understand how neurons in each population responded to the visual stimulus relative to the saccade, a visuomotor index (VMI) was calculated using the formula below (Bruce & Goldberg 1985, Lawrence et al 2005, Sato & Schall 2003, Sommer & Wurtz 2000):

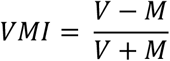

Where *V* is the response in the visual window and *M* is the response in the saccade window, with no baseline subtraction. Therefore, a VMI corresponding to 1 indicates a response exclusively for the visual stimulus, −1 exclusively for the saccade, and 0 indicates equal responses for the visual stimulus and saccade. For FEF, VMI was calculated individually for the 8 conditions and then averaged across conditions to produce a single VMI value per neuron. For 8Ar, a subset of conditions (8 of the 40 conditions at a single amplitude) which most closely approximated the amplitude and direction of the FEF stimuli were selected to facilitate comparison between the two areas.

##### Cross-temporal decoding analysis

A Poisson Naïve Bayesian decoder was implemented to determine the working memory signal readout of 8Ar and FEF populations. For both regions, a pseudo-population was created by combining neurons across recording sessions. Trial to trial dependencies within a session were removed by shuffling the order of the repeats. Any recording session with less than 40 repeats of each condition (1 session for FEF, 4 sessions for 8Ar), and any units that did not have an SNR greater than 2.5 or fire at least 1 spike per second during the delay period of at least one condition were omitted (leading to the removal of 93 FEF units and 561 8Ar units). The instantaneous firing rate of each neuron (100 ms overlapping windows stepped by 50 ms) was used to build a decoder to predict the 8 saccade directions in FEF, and the 8 saccade directions closest in amplitude to the FEF saccade directions for 8Ar (8 of the 40 conditions). The training data set contained 80% of the trials, creating a Poisson distribution model for each direction (θ) using the average spike count for each unit (n_spike_) in the time epoch specified. The remaining 20% of trials were used for testing, at time windows beginning at fixation and ending after the saccade. For a given test trial, the direction with the maximum prediction probability, P(θ| n_spike_), was defined as the predicted saccade direction. P(n_spike_| θ), was calculated using the Poisson distribution model that resulted from the training data. We used 5-fold cross validation, rotating the training and testing data such that each trial was used once for testing, with the average decoding accuracy computed across folds.

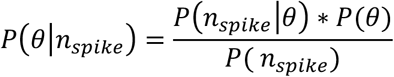

##### Comparison of decoding between 8Ar and FEF

To compare overall decoding accuracy between FEF and 8Ar, we randomly selected a single set of 8Ar neurons (770 neurons) from the total 8Ar population (1722 neurons) to match the size of the recorded population in FEF. When comparing decoding accuracy as a function of other properties (number of neurons, directional selectivity, and reliability) we used the training and testing time point that had the highest accuracy for each area during the delay period (FEF: 50 ms to 150 ms after stimulus offset, 8Ar: 100 ms to 200 ms after stimulus offset).

##### Decoding accuracy and reliability

We developed an index of the reliability of a neuron by calculating a tuning curve separately for the even and odd trials in the time bin with the highest decoding accuracy (see *Methods* above). The reliability was calculated as the Pearson correlation coefficient of the even and odd trial tuning curves. A neuron with an identical tuning curve on even and odd trials would have a reliability of 1, while a neuron in which the tuning curves on the even and odd trials were independent would have a reliability of 0 (on average). Reliability values less than 0 could occur by chance, but would not be expected on average because it would require the tuning curve to shift systematically in preferred direction between the even and odd trials. Each neuron in the pseudo-population was then sorted according to their reliability. Subpopulations of 100 neurons were used to decode eye movement direction, starting with the 100 neurons with the highest reliability, then the next 100 ranked neurons in non-overlapping bins until the remaining population did not have 100 neurons. For the FEF population, this resulted in 7 bins (ranked neurons 1-100, 101-200, 201-300, 301-400, 401-500, and 501-600, and 601-700). For the 8Ar population, the number of 100-neuron bins was larger due to the larger number of 8Ar neurons recorded.

##### Decoding accuracy and tuning selectivity

The selectivity of each 8Ar and FEF neuron was computed during the time window of maximum decoding accuracy (see *Comparison of decoding between 8Ar and FEF*) using a normalized vector strength metric (Smith et al 2002). To measure the selectivity of each neuron’s tuning curve, we calculated the complex summed response vector (where 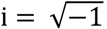)

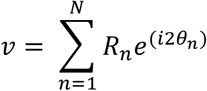

Where R_n_ is the response magnitude during the delay period, θ_n_ is the stimulus location, and n is an index from 1 to the number of points, 8, in the tuning curve. This was then normalized by the summed magnitude of all the response vectors:

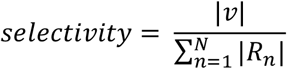

A selectivity of 0 corresponded to a neuron that fired for all conditions equally while a value of 1 indicated a neuron that responded exclusively to one condition. We ranked and grouped neurons based on their selectivity for the decoding analysis in the same manner described above for the reliability analysis.

## Results

We recorded from 2511 8Ar neurons across 39 sessions and 889 FEF neurons across 50 sessions (see *Methods*) in four macaque monkeys while the animals performed a memory guided saccade task (Figure 1B). We sought to understand the principles by which visual and motor signals align and evolve over time during sensorimotor integration. By comparing two brain regions, one closer to the motor output and one further (Leichnetz & Goldberg 1988, Segraves & Goldberg 1987, Sommer & Wurtz 2000), we were able to directly compare the strength and alignment of visual and motor signals from the appearance of a visual stimulus to the execution of a saccade.

### Spatial constancy in 8Ar single neurons

To understand how visual and motor signals are processed at the population level in 8Ar, we first wanted to ensure robust responses were observed at the single neuron level. Previous studies examined visual (Funahashi et al 1989) and motor (Funahashi et al 1991) responses in 8Ar during an oculomotor task, reporting a wide variety of response properties including significant tuning for the visual, delay, and/or saccade epochs, ipsilateral and contralateral tuning, and both excitation and suppression in delay period activity. Having visual stimuli and saccades of numerous amplitudes and directions allowed us to form detailed response fields in the visual and saccade epochs for all neurons recorded. In Figure 2, we show two example 8Ar neurons with large responses during the visual and saccade epochs. Of note, each neuron had a spatially defined area of high firing rate (red) that remained localized to the same region of retinotopic space between visual and saccade epoch – in other words, the tuning was aligned. We observed three key features of 8Ar neuronal responses that are evident in the examples in Figure 2: (1) neurons exhibited both excitation and suppression relative to their baseline rate (particularly evident in Figure 2B), and regions of peak suppression tended to be located 180 degrees away from regions of peak excitation, (2) neurons typically were tuned in their responses smoothly across the whole tested visual field, as opposed to the punctate receptive fields characteristic of early visual cortex, and (3) excitatory response regions could be located either contralateral (Figure 2A) or ipsilateral (Figure 2B) to the recorded hemisphere.

**Figure 2:**
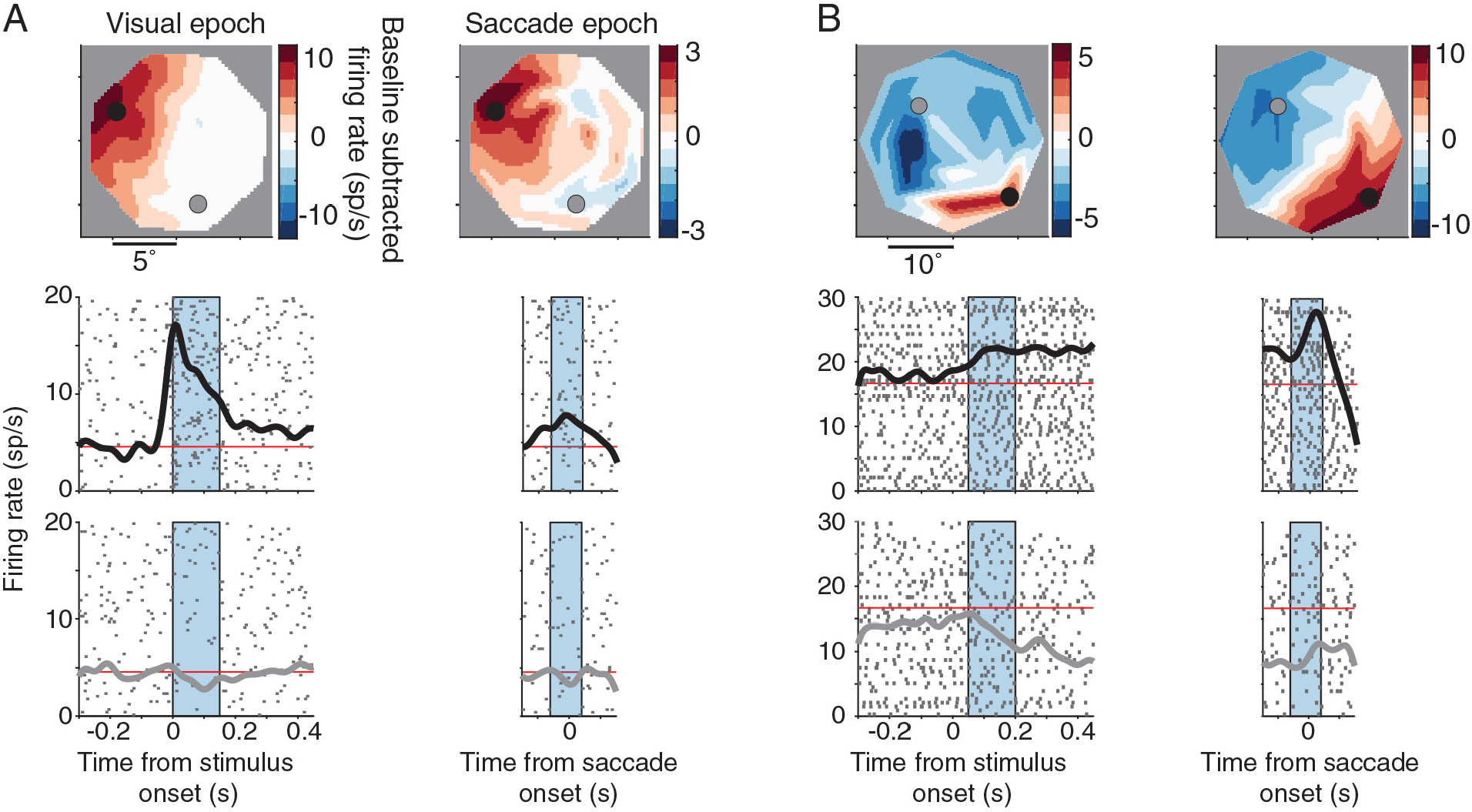
Spatial constancy in 8Ar neurons. A) Top: Response field map for an example neuron during the visual and saccade epoch. Firing rate was baseline subtracted (170 ms to 20 ms before stimulus onset) such that red colors indicate activity above baseline, blue colors activity below baseline, and white near baseline. Middle: PSTHs aligned to stimulus onset and saccade onset for a condition close to the center of the response field (black circle). Bottom: PSTHs aligned to stimulus onset and saccade onset for a condition in the opposite hemifield of the center of the response field (gray circle). The blue shaded regions indicate the time period in which the firing rate was calculated during the visual and saccadic epochs for the response field maps. This neuron had a robust visual and saccadic response that was localized to the contralateral hemifield and was spatially congruent between the visual and saccade epoch. B) An example neuron with a robust and spatially congruent visual and saccadic response localized to the lower portion of the ipsilateral hemifield. Spatial locations opposite the center of the response field were suppressed below baseline. Note: all response field maps were flipped such that the left hemifield represented the contralateral hemifield. Example A was from monkey Wa and example B was from monkey Pe.

### Response latency in FEF and 8Ar

Having confirmed that 8Ar neurons had distinct and spatially localized response fields in both the visual and saccade epochs, we next identified the ideal time window to accurately capture the visual and saccadic responses. Previous studies have demonstrated 8Ar visual responses can have a variety of time courses (Mikami et al 1982, Suzuki & Azuma 1983): transient bursts of excitation or suppression after stimulus onset or perisaccadically, sustained modulation throughout the entire delay period, or a combination thereof. We first examined the population PSTH aligned to the visual stimulus or saccade and measured the overall time course of 8Ar activity. The 8Ar population had a small visual transient with a longer sustained period of activity during the delay period, which then rose perisaccadically and peaked after saccade onset (Figure 3A). This contrasted with the FEF population PSTH, which had a more phasic visual transient at a shorter latency, as well as perisaccadic activity that peaked closer to saccade onset (Figure 3B). To determine the latency of each neuron in response to a visual stimulus and relative to a saccade, we calculated the variance across the target conditions in 50 ms windows (see *Methods*). To ensure a fair comparison between 8Ar and FEF latencies, the same window width and epoch times were used. For neurons with significant visual responses (p < .01, Kruskal-Wallis test), 8Ar had a significantly longer latency when compared to FEF (Figure 3C, 8Ar mean = 189 ms, FEF mean = 154 ms; two sample t-test p < .001), consistent with our visual observations of the PSTHs in the two areas. Our estimate in FEF was later than other reports (such as (Mayo et al 2015, Schmolesky et al 1998), in part because our latency metric measures peak tuning and not response onset as in some other studies. This latency difference, combined with the visual comparison of the PSTHs, suggest a substantially more robust and earlier visual response in FEF than in 8Ar.

**Figure 3:**
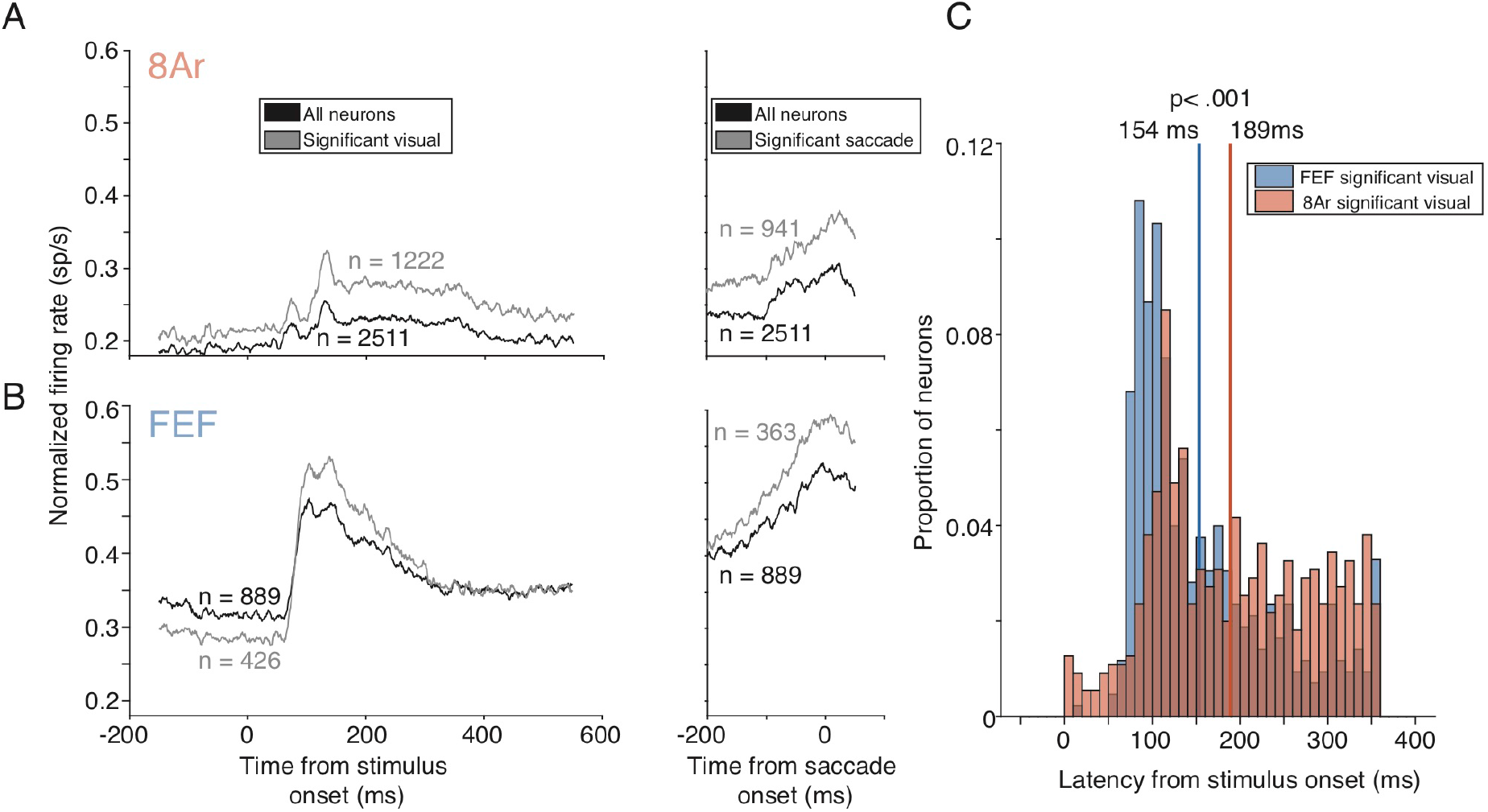
Latency in 8Ar and FEF. A) Population PSTH for all 8Ar neurons (black line, n = 2511 neurons) and significantly tuned 8Ar neurons (grey line; p < .001 Kruskal Wallis test) in the visual (left) or saccade (right) epoch. Significant visual neurons (n = 1222 neurons) passed the significance test in the visual epoch while significant saccade neurons (n = 941 neurons) passed the test in the saccade epoch. Each neuron’s PSTH was normalized by the maximum firing rate in either the visual/delay epoch (0 ms to 550 ms after stimulus onset) or the saccade epoch (−200 ms to 50 ms before saccade onset). B) Same convention as in A, but with all FEF neurons (black line; n = 889 neurons) and significantly tuned neurons (grey line) in the visual (n = 426 neurons) or saccade (n = 363 neurons) epoch. The 8Ar population PSTH had less modulation with respect to baseline and a longer latency compared to FEF in the visual and saccade epochs. C) Distribution of single neuron latencies for FEF (blue) and 8Ar (orange) during the visual epoch. Only neurons with significant visual responses were included (8Ar = 1222 neurons; FEF = 426 neurons). The 8Ar distribution had a significantly longer visual latency compared to FEF (p < .001; two sample t-test).

### Hemifield tuning differences in 8Ar

Previous 8Ar studies have found neurons tuned to stimuli in the ipsilateral visual field (Funahashi et al 1989, Funahashi et al 1991), unlike most earlier visual areas which are entirely contralateral or extend only minimally into the ipsilateral hemifield (Gattass et al 1981, Gattass et al 1988). Our observations of individual example neurons (Figure 2) extend beyond a simple classification of contralateral or ipsilateral – individual neurons demonstrated smoothly varying responses across the whole tested visual field. We sought to determine whether this observation in individual neurons was representative of the entire population, and then asked how the ipsilateral and contralateral representations of space differ in 8Ar neurons.

We first defined a neuron as ipsilateral or contralateral based on the location of the center of mass calculated on the response that was elevated above baseline (the red region of the response maps, see *Methods*). For each group (ipsilateral and contralateral), we rotated the response field map of each neuron such that its center of mass was on the horizontal meridian. Ipsilateral and contralateral tuned neurons were then combined separately to form a population response field map for the visual epoch and the saccade epoch. For the contralateral tuned population, suppression in the opposite hemifield was more apparent in the saccade epoch compared to the visual epoch (Figure 4A). For the ipsilateral tuned population, suppression was equally present in both visual and saccade epochs (Figure 4B). The observation of stronger suppression in the ipsilateral tuned population compared to the contralateral tuned population during the visual epoch agrees with a previous finding (Bullock et al 2017).

**Figure 4:**
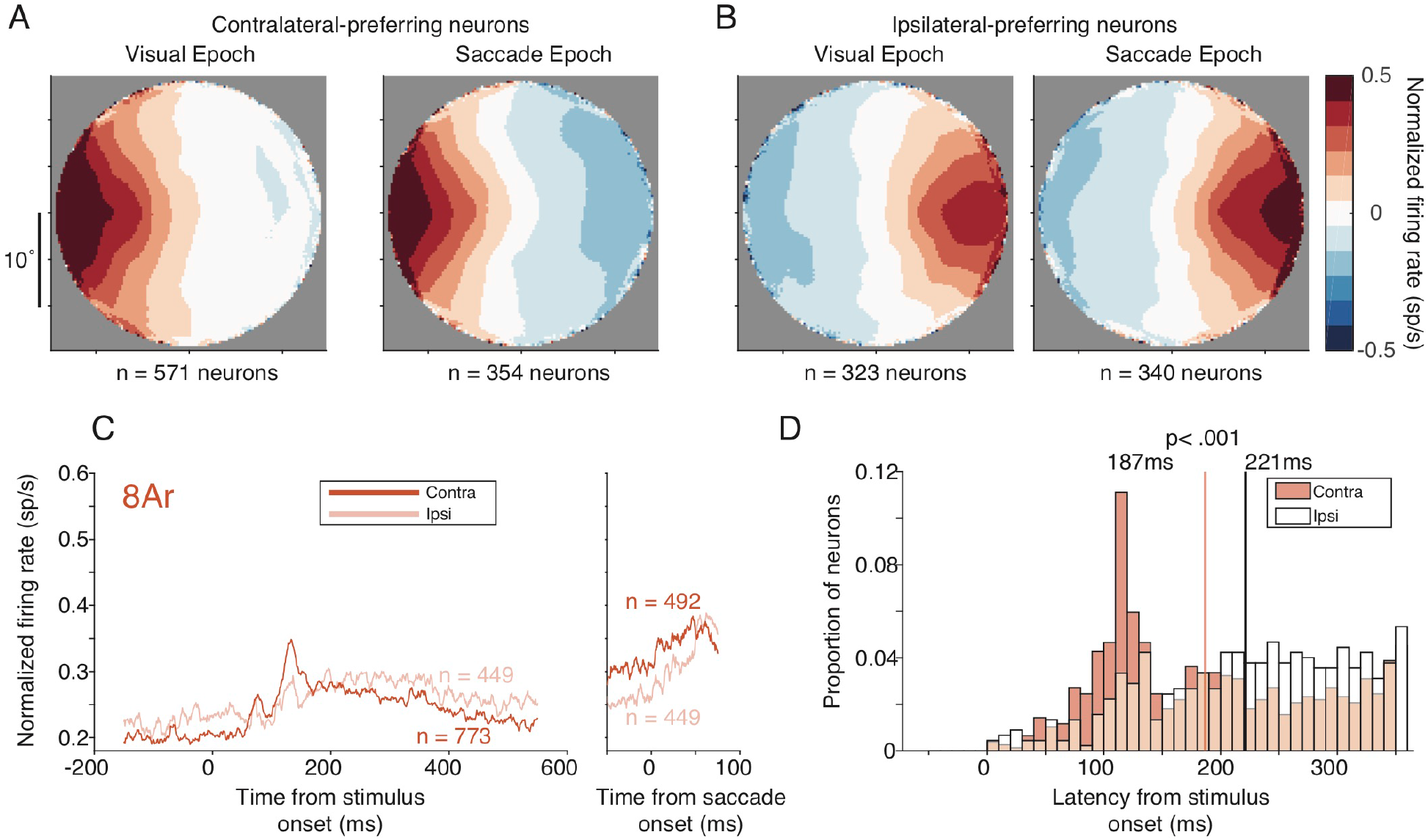
Hemifield tuning differences in 8Ar. A) Population response field maps for contralateral tuned neurons in the visual (left) and saccade (right) epoch. Each neuron’s response field map was normalized by the maximum response and rotated to the horizontal meridian. All normalized and rotated maps were then averaged across neurons to yield the population response field map. B) same convention as in A but for the ipsilateral tuned neurons. The ipsilateral and contralateral populations had similar suppression during the saccade epoch, however less suppression was observed for the contralateral population during the visual epoch. C) Population PSTHs for contralateral (dark orange) and ipsilateral (light orange) neurons aligned to stimulus onset or saccade onset. During the visual epoch, the ipsilateral population PSTH had less modulation relative to baseline and a later latency than the contralateral population. During the saccade epoch, the ipsilateral population PSTH had more modulation relative to baseline but still had a longer latency. D) Distribution of latencies for contralateral (filled) and ipsilateral (open) neurons during the visual epoch. Only neurons with significant visual responses were included (contralateral = 773 neurons; ipsilateral = 449 neurons). The ipsilateral distribution had a significantly longer visual latency compared to the contralateral distribution (p < .001; two sample t-test).

We then investigated the contralaterally and ipsilaterally tuned population PSTHs aligned to stimulus and saccade onset. For the visual epoch, the ipsilateral tuned population had a later and weaker visual transient, coupled with a stronger sustained level of activity in the delay period after the visual transient (Figure 4C). During the saccade epoch the ipsilateral population began at a lower level of activity and increased more sharply perisaccadically. Comparing the visual latency distributions, ipsilateral neurons had significantly longer visual latencies (p < .001, two sample t-test) and had a more uniform distribution of latencies compared to the contralateral distribution, which had a clear peak around 187 ms (Figure 4D).

### Dynamic selectivity in 8Ar

So far, we have demonstrated 8Ar neurons had a wide variety of spatial tuning and latencies across the visual and saccade epochs. Given this variety, we sought to understand how these visual and motor signals coexisted within 8Ar. One particularly intriguing aspect of dynamic selectivity in 8Ar, observed at the single neuron level, is an alteration of the spatial response preferences during the delay period of a working memory task, often referred to as mixed selectivity (Parthasarathy et al 2017, Spaak et al 2017). However, typical investigations of this property employed a limited set of conditions, displaying stimuli at only one eccentricity. Our experimental paradigm tiled a substantially larger portion of visual space resulting in a more detailed estimate of each neuron’s visual and saccadic response field. Using this high resolution, we sought to confirm the extent of dynamic selectivity across the population of 8Ar neurons and determine if there were systematic rules by which this representation evolved over the time period between the visual stimulus and the saccade.

We found that so-called mixed selectivity existed in a subset of 8Ar neurons, and comparing single neuron examples illuminated subtle differences in how individual neurons changed their tuning. For some neurons, the center of the response field shifted drastically between the visual and motor epochs, such as a 90-degree rotation (Figure 5A). For other neurons, the response field during the visual epoch broadened during the saccade epoch, such that stimuli that were suppressed during the visual epoch became regions of peak activity during the saccade epoch (Figure 5B). Finally, some neurons shifted the center of their response field 180-degrees, where the area of maximum activity during the visual epoch was suppressed below baseline during the saccade epoch (Figure 5C). These results provide clear examples of single neurons shifting their tuning preferences between the visual and saccade epochs and highlight the diversity of spatial shifts observed across individual neurons.

**Figure 5:**
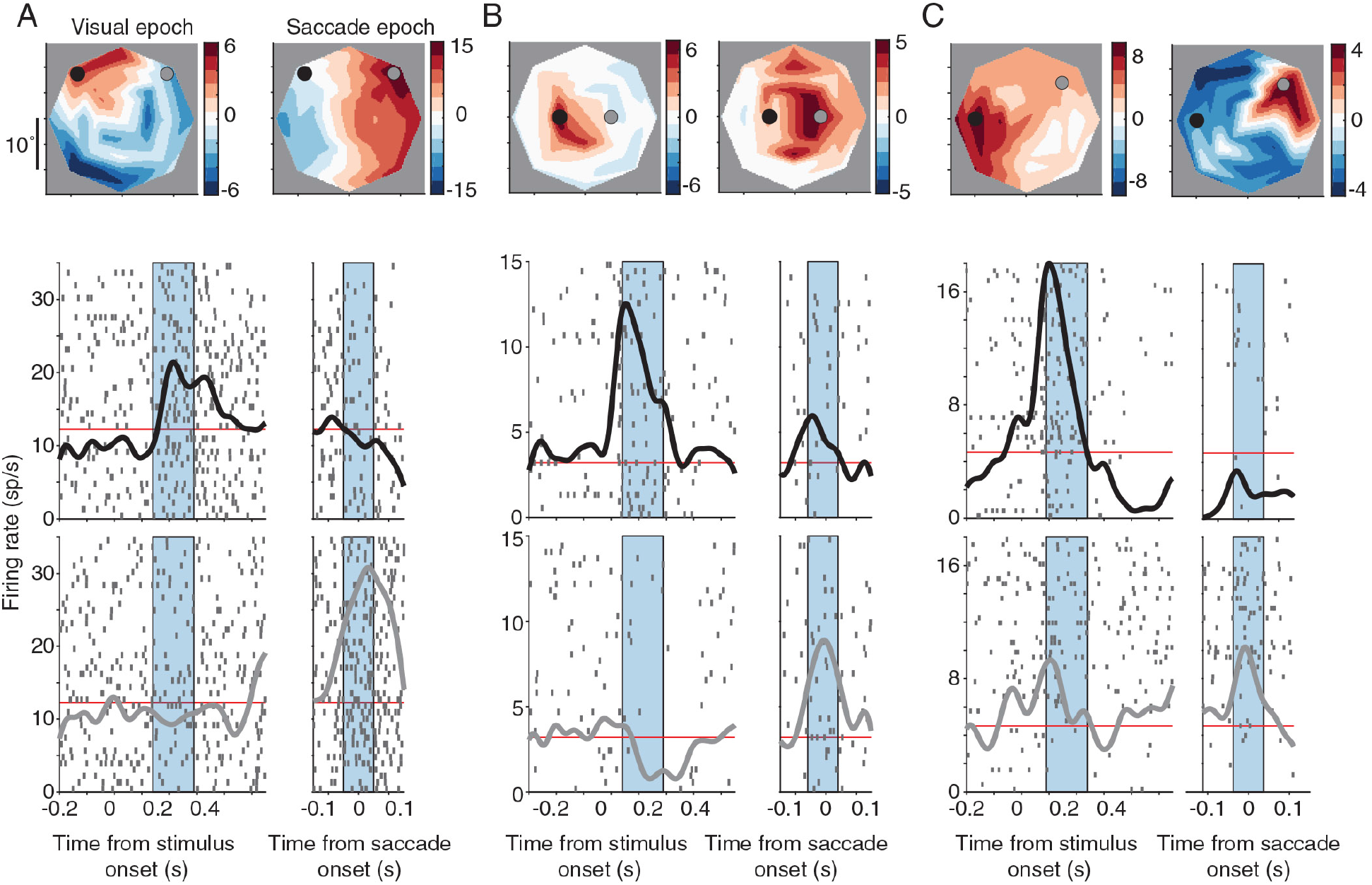
Dynamic selectivity in single neurons. A) Top: response field map of the baseline subtracted firing rate for an example neuron. Bottom: Average PSTH for a condition close to the center of the visual response field (black) and close to the center of the saccade response field (grey). This example neuron was contralaterally tuned during the visual epoch but rotated its response field 90-degrees to the ipsilateral hemifield during the saccade epoch. B) Example neuron with contralateral tuning during the visual epoch, with the ipsilateral hemifield suppressed. During the saccade epoch, the response field broadened such that the neuron fired above baseline for the condition that was previously suppressed. C) Example neuron with a robust visual response in the contralateral hemifield that is suppressed during the saccade epoch. The response field shifted nearly 180-degrees between the visual and saccade epochs. Examples A and B were from monkey Pe, example C from monkey Wa.

The previous examples of mixed selectivity compared the tuning of individual neurons in specific windows during the visual and saccade epochs. If the visual stimulus and saccade signals are indeed separately coded in 8Ar, and the switch from a visual to a saccadic code accounted for much of the diversity seen in the 8Ar neuronal response, we hypothesized the maximum difference in the response field of the visual and saccade epochs should occur between the peak of the visual response and the onset of the saccade. To determine whether this was the case, we calculated the center of the response field in 50 ms windows throughout the entire trial (fixation onset to saccade onset). To understand how the center of the response field shifted over the course of the trial, we subtracted the angle associated with the center of the response field in the ideal visual epoch window (see *Methods*) from the angle associated with the center of the response field calculated in sliding windows throughout the trial. If a neuron maintained the spatial location of its visual response field throughout the entire trial, subtracting by the preferred location would yield an angular difference of 0 throughout the trial (Figure 6A, top). Conversely, if a neuron shifted its tuning during the saccade epoch, we would predict the angular difference to be low in the visual epoch (as it is close to the ideal visual time window) but increase as the time window approached the saccade (Figure 6A, bottom). The same process was repeated for the ideal saccade window, where the center of the response field throughout the trial was subtracted by the ideal saccade window. These angular difference curves throughout the trial were calculated for each neuron and then averaged across neurons to yield a population metric. We found the maximum angular difference between the center of mass at a given time in the trial and the ideal visual window corresponded to saccade onset, while the minimum angular difference was observed around the mean visual latency of the population (180ms after stimulus onset) (Figure 6B, left). Conversely, the maximum angular difference for the ideal saccade window was after stimulus onset, and the minimum angular difference was at saccade onset (Figure 6B, right). Thus, the spatial shifts in 8Ar neurons were most obvious when comparing the visual and saccade-aligned responses.

**Figure 6:**
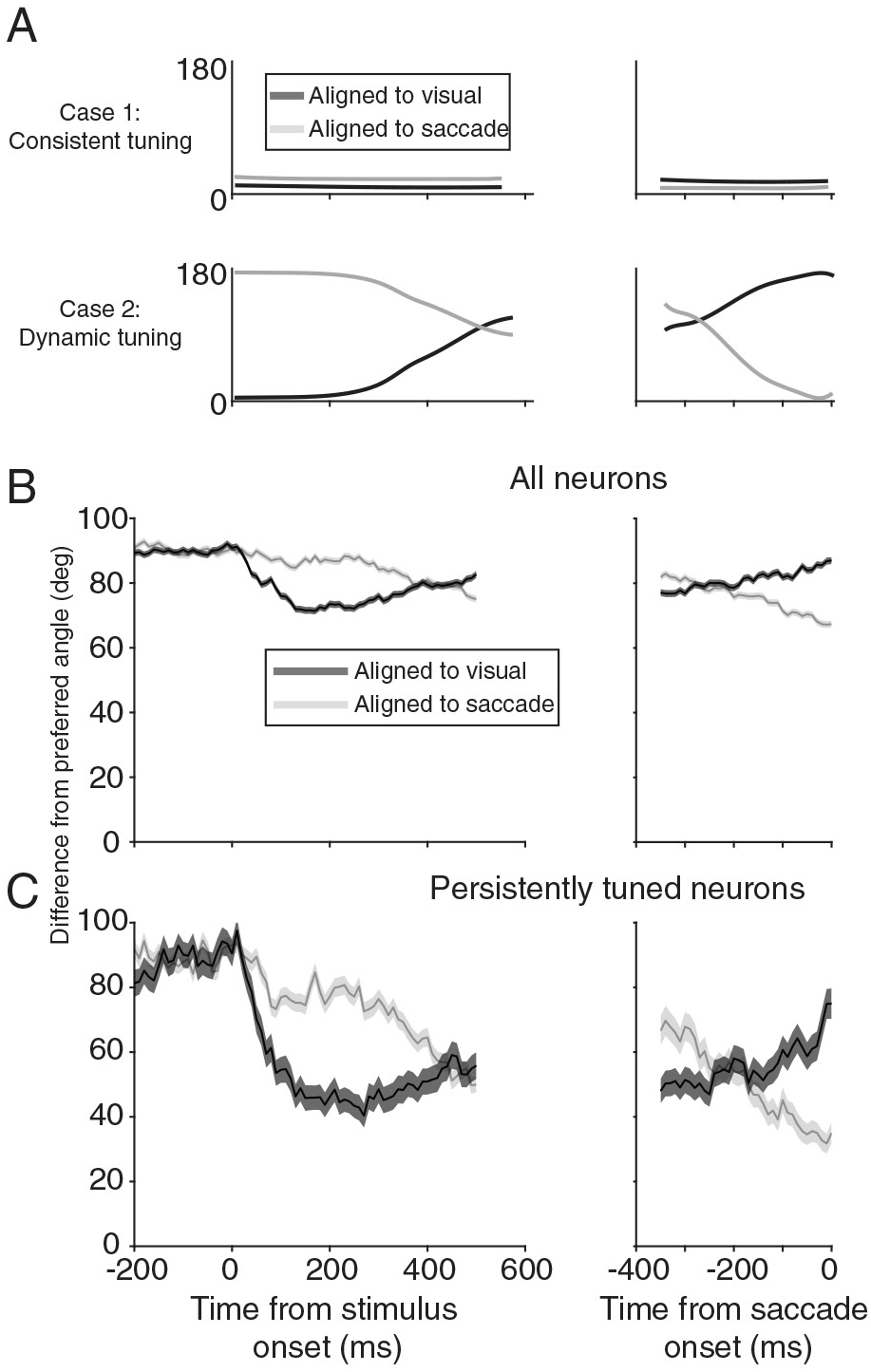
Time course of visual and motor selectivity. A) Illustration of the angular difference expected throughout the time course of the trial for an idealized neuron with consistent tuning (top) and dynamic tuning (bottom) between the visual and saccade epochs (with a zero-latency visual response). B) Angular difference for all 8Ar neurons (n = 2511 neurons) between the center of the response field at a specific time during the trial and the center of the response field for the ideal latency in the visual (black) or saccade (grey) epoch aligned to stimulus (left) or saccade (right) onset. C) same convention as in B however only for a subpopulation of neurons that had persistent significant tuning during the visual and saccade epochs (n = 158 neurons). The time during the trial at which the response field center was furthest from the response field calculated during the ideal visual latency was saccade onset. Conversely, the time during the trial at which the response field center was furthest from the response field calculated during the ideal saccade latency was in the 200ms following stimulus onset.

Large shifts in the center of the response field could be confounded by neurons that did not fire in the other epoch, thus creating a noisy estimate of the center of the response field. To address this, we selected 8Ar neurons which were tuned both to the location of the visual stimulus as well as the saccade. We found those neurons in the population which had significant tuning (Kruskal Wallis test, p < .001) in at least 50% of the time points following the visual stimulus (50 ms to 350 ms after stimulus onset) and preceding the saccade (−100 ms to 0 ms before saccade onset). The rationale was these neurons maintained their tuning in the visual and saccade epochs, and thus any shifts in tuning were not due to neurons that had a strong spatial preference in one epoch but weak or noisy responses in the other. Within this subpopulation, the same angular difference trends were maintained (Figure 6C). In summary, the maximum shift in response fields occurred between the visual and saccade epochs and this shift was not confounded by neurons that were untuned in either of the epochs. This is consistent with the hypothesis that some of the rich dynamics observed in 8Ar emerge due to a transition between separate visual and motor representations in the population.

### Comparison between FEF and 8Ar

Based on the results presented up to this point, it is clear a subpopulation of 8Ar neurons altered their tuning between the visual and saccade epochs even though the visual stimulus and saccade endpoint were at the same spatial location. Furthermore, this was not simply due to a loss of tuning during one of the epochs. If mixed selectivity occurred in 8Ar, a natural first step would be to determine whether this property was unique to a subpopulation of 8Ar neurons, such as the ipsilateral neurons that we found had longer response latencies and different patterns of suppression opposite the response field. In addition, we considered whether other cortical regions exhibited similar changes in tuning, or whether this property was unique to 8Ar. To determine the relative prominence of neurons with shifting tuning in 8Ar, we compared our observations at a population level with FEF. Given FEF is more closely linked to the generation of saccades, we hypothesized more FEF neurons would have a congruent alignment of their visual and motor signals compared to 8Ar.

We first asked whether there was any pattern to how 8Ar neurons shifted their response profiles between the visual and saccade-related responses. To do this, we included only neurons that were selective in both the visual and saccade epochs (p < .001; Kruskal Wallis Test; Contra: n = 412 neurons; Ipsi: n = 217 neurons). We then computed the angular difference between the center of the response field in the ideal visual and saccade epoch and binned these angular differences in six 30° bins. We compared the distribution of these visual-motor angular differences to a null distribution obtained by associating each neuron’s preferred visual response angle with the preferred saccadic response angle of a different neuron. We repeated this process 1000 times to obtain the 5^th^ and 95^th^ confidence intervals for each 30° bin. For both the contralateral and ipsilateral populations in 8Ar, we found that roughly half of the neurons had visual and saccadic peak response angles within 60° (Figure 7A). Of the remaining neurons, there was a tendency for a mirror inversion (> 120° shift) more often that an intermediate rotation (60-120°).

**Figure 7:**
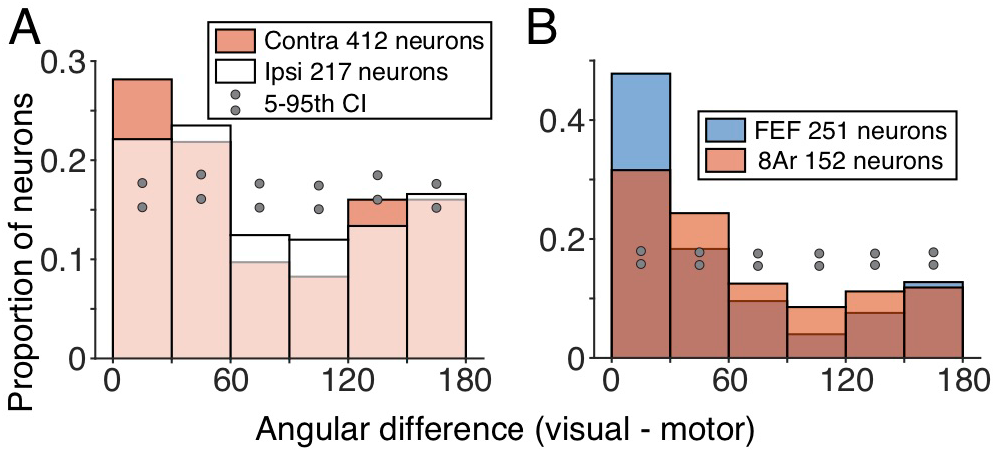
Tuning shifts in 8Ar and FEF subpopulations. A) Shifts in response field centers between the visual and saccade epoch as measured by angular difference for contralateral (filled) and ipsilateral (open) tuned neurons. Only neurons with significant visual and saccade responses were analyzed (p<.01, Kruskal Wallis Test; Contra n = 412; Ipsi n = 217). 5^th^ and 95^th^ percent confidence intervals show the proportion of neurons expected for a given angular difference by chance (upper and lower grey dots). Contralateral and ipsilateral populations had similar distributions with respect to angular difference. Many neurons had consistent visual and saccade alignment, however a substantial portion also shifted their tuning dramatically. B) same conventions as in A but comparing the FEF population to the 8Ar population. Some FEF neurons had substantial shifts in tuning, equivalent to the 8Ar population, however FEF also had more neurons with congruent visual and motor response fields.

To compare the shifts we observed in 8Ar with a baseline from an area that has been studied extensively in visuomotor tasks, we matched our data in the two areas across conditions and trial repeats (see *Methods)*. This meant a subset of the 8Ar data were used (1 target amplitude, 8 directions), and all FEF and 8Ar sessions were randomly subsampled to 40 trial repeats per condition. The same method for calculating latency was used on this subsampled data from both areas to calculate the ideal visual and saccade response window and to identify significantly tuned neurons (p < .001; Kruskal Wallis test; FEF n = 251 neurons, 8Ar n = 152 neurons). Our results in 8Ar with this reduced data set were similar to those obtained in the full 40-condition data (Figure 7B). As a population, FEF had a greater proportion of well-aligned neurons (< 30° angular difference) than 8Ar (8Ar = 32%; FEF= 48%; p = .0014, Chi-square test). However, both areas contained a subset of neurons that reliably shifted their tuning between the visual and saccadic epochs.

### Visual and motor response properties in FEF and 8Ar

In addition to the spatial response profile, we sought to compare the relative strength of visual and motor signals in 8Ar and FEF by computing a visuomotor index (VMI, see *Methods*) across conditions for each neuron in the FEF and 8Ar populations. Broadly, these distributions were quite similar in the two regions – most neurons exhibited some degree of visual and saccadic responses, leading to distributions centered on zero. However, relative to the FEF distribution, 8Ar was significantly (p = .003; two sample t-test) shifted toward 1, meaning 8Ar neurons were more likely to have a stronger visual response compared to a saccade response (Figure 8A). In addition to this difference in the ratio of visual and saccadic responses, FEF had a larger proportion of significantly tuned neurons (p < .001 in the visual, motor, or both epochs; Kruskal Wallis test) in visual, visuomotor, and motor groups (chi squared test; visual: p = .01; motor: p = .007; visuomotor: p < .001) (Figure 8B). When combining the three groups (tuned visual, motor, and visuomotor) FEF also had a significantly higher proportion of contralaterally tuned neurons (chi squared test, p = .002; FEF 68% contra, 8Ar 59% contra). The presence of more ipsilateral tuned neurons in 8Ar was even more striking when considering only the most directionally selective (see *Methods*) neurons in each population (Figure 8C). To obtain these selective neurons for a given visual/motor/visuomotor group, a neuron had to be significantly tuned (p < .001; Kruskal Wallis test in one or both of the visual and saccadic epochs) as well as have a directional selectivity (see Methods) that was at or above the 90^th^ percentile for the epoch. For most of the groups in both 8Ar and FEF, selective neurons had a strong contralateral bias (FEF contralateral: visual 82%, motor 71%, visuomotor 79%; 8Ar contralateral: visual 79%, motor 30%, visuomotor 71%). In FEF, this result is consistent with a previous study by our group that found almost exclusively contralateral RFs in a population of neurons with brisk visual responses to a dynamic dot stimulus (Mayo et al 2015). Interestingly, when considering only these selective neurons, the 8Ar motor population had more ipsilateral than contralateral neurons. Upon visual inspection of these ipsilateral tuned motor neurons, we noted many of them had drastic shifts in their spatial responses between the visual and motor epochs, with contralateral tuned visual responses (that did not pass the significance test for tuning in the visual epoch). Overall, these results indicate that FEF neurons are more strongly tuned, and more contralaterally biased, than 8Ar neurons, which have a more balanced representation of space that also appears more likely to shift between the visual and motor epochs.

**Figure 8:**
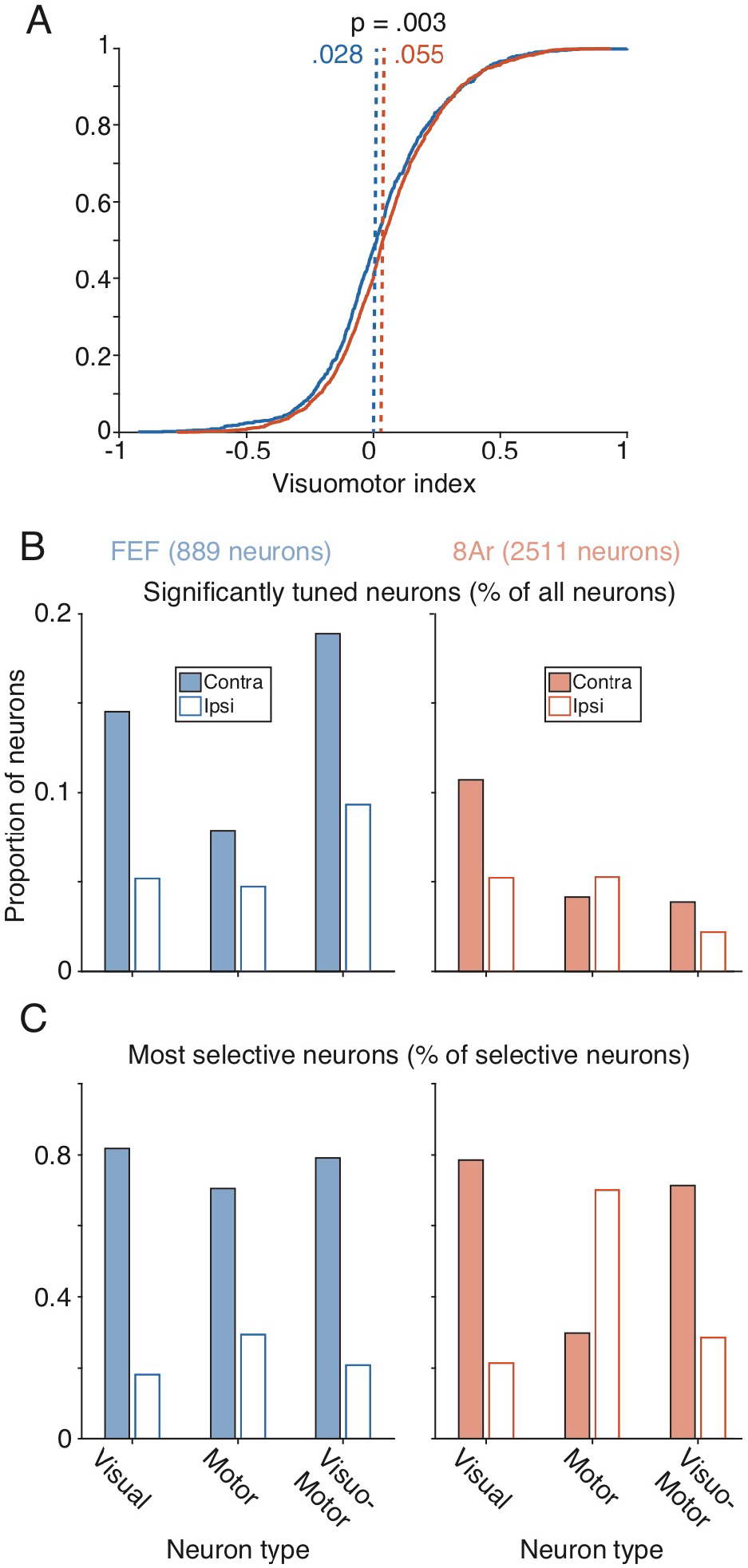
Visual/motor tuning in 8Ar and FEF. A) Cumulative distribution of visuomotor index for 8Ar and FEF populations, all neurons included (FEF n = 889 neurons; 8Ar n = 2511 neurons). 8Ar neurons had a stronger visual response when compared to the saccadic response. B) Distribution of visual, motor, and visuomotor neurons within FEF (left) and 8Ar (right), normalized to the total number of neurons recorded. Groups were divided by spatial tuning (contralateral: filled, ipsilateral: open). To be included a neuron needed to have significant tuning in at least one epoch or both (visuomotor group). FEF had more tuned neurons for all groups (visual, motor, visuomotor). C) Same convention as in B, but with the additional criterion that each neuron had to be in the 90^th^ percentile or above ranked by directional selectivity (FEF: visual = 22, motor = 17, visuomotor = 24; 8Ar: visual = 70, motor = 57, visuomotor =12). A majority of the most selective FEF neurons had contralateral tuning, while in 8Ar, highly selective motor neurons were more likely to be ipsilateral tuned.

### Decoding from 8Ar and FEF populations

Individual 8Ar neurons had numerous tendencies consistent with the mixed or dynamic selectivity observed by other groups during working memory tasks. One approach to quantify the effect of these changes in individual neurons is to apply decoding analyses to the whole population (Astrand et al 2014, Barak et al 2010, Parthasarathy et al 2017, Spaak et al 2017, Stokes et al 2013). An accumulation of tuning shifts across individual neurons would lead to a spatial representation that did not generalize well across time. In such a situation, a decoder built on data from one time point in the trial would do poorly in predicting target location at another time point. Our results with individual neuron analyses in 8Ar and FEF led us to predict that the population-level signal in FEF would be more temporally generalizable than that in 8Ar.

We used a Poisson Naïve Bayes decoder, trained on neural activity from one time window in the trial, and tested on all other time points during the trial (see *Methods*). We combined neurons across recording sessions to create a pseudo-population in FEF and 8Ar, using 8 conditions (1 amplitude, 8 directions) and, to normalize across sessions, 40 trial repeats randomly selected from each condition with the trial ordering shuffled for each neuron to destroy any correlations between neurons. All decoding accuracies reported were the average across the eight conditions, and standard errors were computed across cross-validation folds. For comparisons between FEF and 8Ar, the 8Ar pseudo-population was randomly subsampled to match the number of neurons in the FEF pseudo-population, unless stated otherwise.

Overall decoding performance, as well as generalizability across time, was higher in the FEF pseudo-population compared to 8Ar. For FEF, decoding was highest shortly after visual onset and around the time of the saccade, but also maintained a high accuracy throughout the delay period (Figure 9A). The generalizability of the FEF population code was seen by examining bins on the off-diagonal, where the training and testing epochs were temporally separated. The decoder trained using FEF activity was more generalizable when compared to 8Ar (Figure 9B). We took a cross-section of the decoding performance map and evaluated the testing accuracy across the trial when the training bin was held constant (Figure 9, inset line graphs). If activity in a given training bin was generalizable across time, the resulting accuracy curves would be broad, while non-generalizable activity would have a sharp peak corresponding to when the training and testing bins temporally aligned. The accuracy curves for 3 training bins throughout the delay period were higher and broader for FEF compared to 8Ar, suggesting the FEF population code had a more accurate readout of the stimulus/saccade encoded during a trial, and that the code was more generalizable throughout the trial.

**Figure 9:**
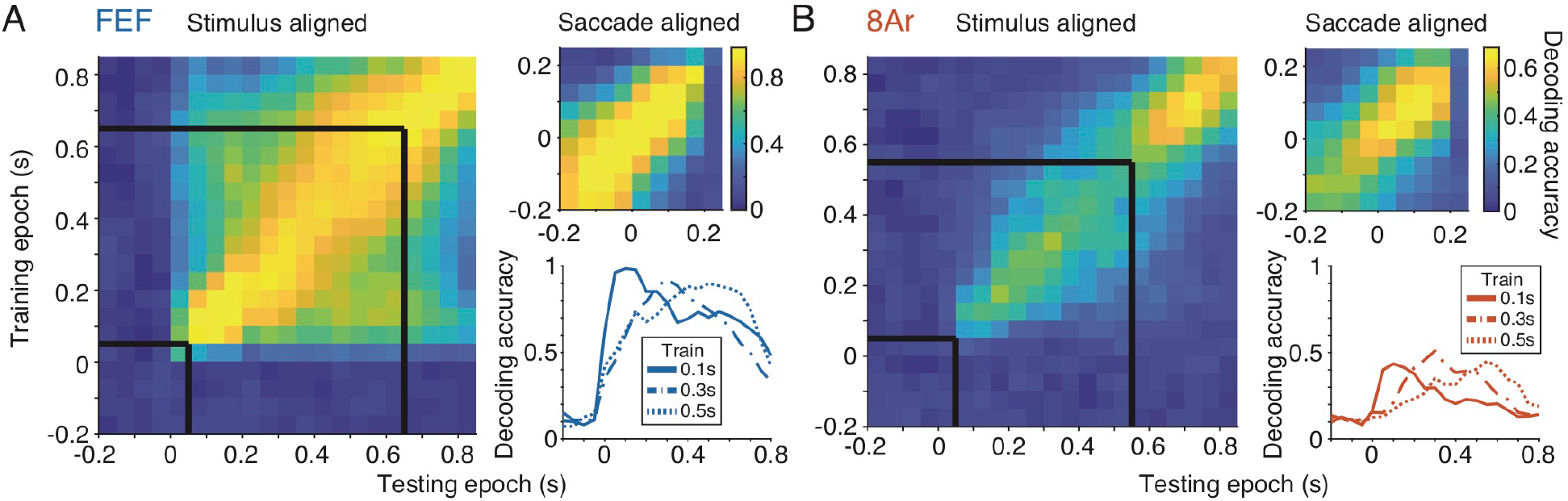
Decoding in 8Ar and FEF. A) Decoding performance of the FEF pseudo-population (n = 770 neurons) for various training and testing points throughout the trial, aligned to stimulus onset or saccade onset (inset). Black lines denote the beginning and end of the delay epoch. B) same convention as in A, but with a 8Ar pseudo-population randomly subsampled to have the same size as the FEF pseudo-population (n = 770 neurons of 1722 neurons).Inset) cross-sections of decoding accuracy, where the training bin was fixed to one of three points during the delay period (0.1, 0.3, or 0.5 seconds after stimulus offset) and the testing bins varied across the entire trial. Decoding accuracy was highest for training and testing points that were temporally in the same bin, particularly after stimulus onset and around the time of the saccade. FEF had a higher overall decoding accuracy and a higher accuracy for training and testing points that were temporally separated.

To understand why the FEF decoder performed better than the 8Ar decoder, we related decoding accuracy to three basic properties of the pseudo-population: the number of neurons in the population, the direction selectivity of the neurons, and their reliability in response (the correlation in tuning curves between even and odd trials, see *Methods*). For this analysis, one time point with the highest decoding accuracy after stimulus onset was selected for testing and training (FEF: 100-150 ms after stimulus onset, 8Ar 150-200 ms after stimulus onset). We first examined decoding accuracy as a function of the number of neurons in the pseudo-population. Across all pseudo-population sizes, the FEF decoder performed better than the 8Ar decoder (Figure 10A). Starting with a population of 100 neurons, the FEF decoder increased in accuracy as more neurons were added and began to asymptote at 100% accuracy for populations over 500 neurons. The 8Ar population started at an overall lower accuracy level and monotonically increased as more neurons were added, but the decoder never reached the accuracy of even the smallest population of FEF neurons we tested (FEF accuracy, 100 neurons 71.5%; 8Ar accuracy 1722 neurons 63.5%).

**Figure 10:**
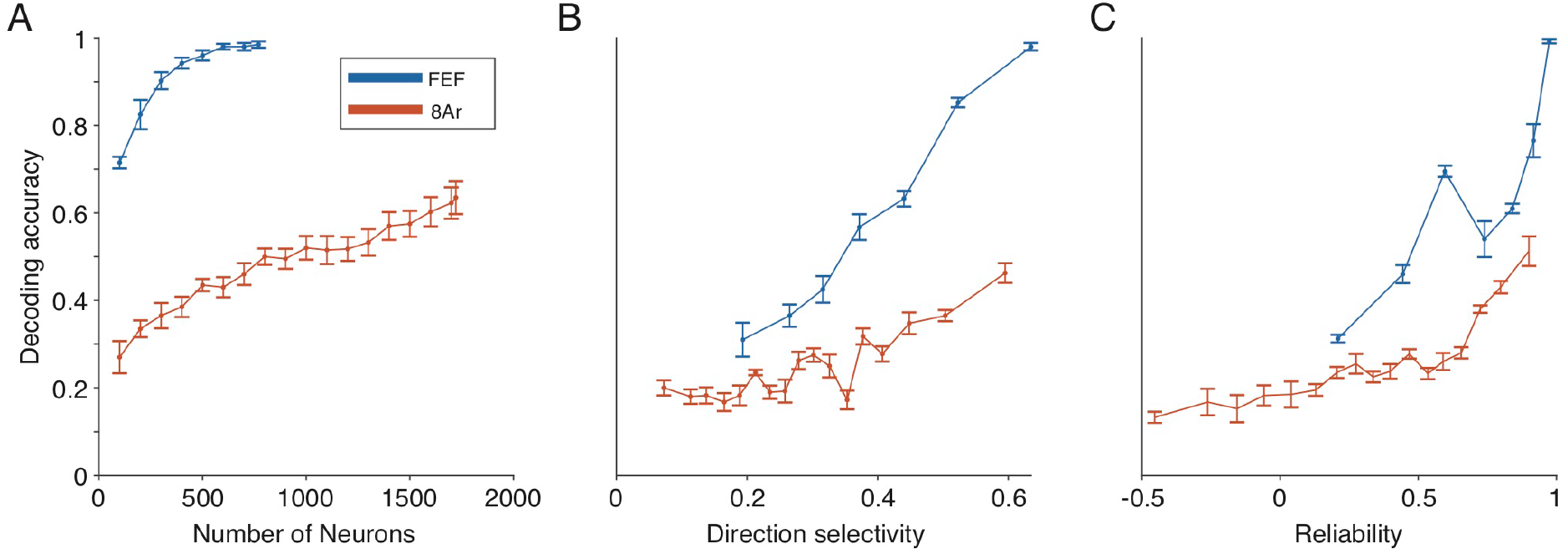
Decoding accuracy with single neuron properties. A) Decoding accuracy in FEF (blue) and 8Ar (red) as a function of the size of the pseudo-population. Even the smallest FEF population tested (100 neurons) performed better than the entire sample of 8Ar neurons (1722 neurons). B) Decoding accuracy for groups of 100 neurons (non-overlapping) as a function of direction selectivity. For both FEF and 8Ar, as the mean direction selectivity of the population increased, the decoding accuracy increased. The difference in decoding accuracy between 8Ar and FEF neurons decreased when matched for direction selectivity, but the FEF populations still maintained a higher decoding accuracy. C) Same convention as in B but matched for reliability. Similar to the direction selectivity results, decoding accuracy increased as reliability increased, and the difference in decoding accuracy between FEF and 8Ar was reduced when matched for reliability, but the decoding performance in FEF remained higher than 8Ar.

Knowing that a decoder trained on activity from a small population of FEF neurons (100 neurons) could outperform one trained on the entire 8Ar population (1722 neurons) we examined what individual response properties could lead to such a wide margin in decoding. We first examined direction selectivity, a measure of tuning across the 8 conditions (see *Methods*). One possibility is that the neurons in FEF were merely more selective, and therefore the population of FEF neurons produced better decoding. For each pseudo-population (FEF and 8Ar), each neuron was ranked by their selectivity. Then, nonoverlapping groups of 100 neurons were chosen starting with the most selective. Decoding accuracy increased with the directional selectivity of the neurons for both FEF and 8Ar. When we compared subpopulations where the average selectivity of the 100 neurons was the same, FEF decoding accuracy was still larger than 8Ar (Figure 10B). However, the much smaller difference in decoding between FEF and 8Ar in matched selectivity groups indicates that one reason for the better decoding in FEF was that FEF neurons were, on average, more selective than 8Ar.

We considered a second response property that could influence decoding, which was the reliability of the neurons from trial to trial for the same stimuli (see *Methods*). Using the same methodology as for selectivity, we ranked the neurons and measured decoding in groups of 100. As with selectivity, decoding accuracy increased with reliability for both FEF and 8Ar populations. In groups of neurons matched by their reliability, FEF decoding performance was closer to, yet still slightly exceeding 8Ar performance (Figure 10C). Taken together, these analyses show that the overall higher selectivity and reliability in FEF neurons are important contributors to the higher ability to decode from small populations in FEF compared to 8Ar.

## Discussion

The transition from perception of a visual stimulus to the generation of a saccadic eye movement is fundamental to primate behavior and has been an important model system for studying the broader process of sensorimotor integration. Using a memory guided saccade task that separates responses due to the visual stimulus from those associated with the eye movement, we studied the dynamics of high-resolution visual and motor representations in 8Ar and FEF. These two cortical regions have been implicated as important in sensorimotor transformations, particularly in the context of spatial working memory, with FEF situated closer to the motor output, with its direct connections to the superior colliculus (Leichnetz et al 1981, Segraves & Goldberg 1987, Sommer & Wurtz 2000) and brain stem oculomotor nuclei (Huerta et al 1986, Leichnetz et al 1984), and 8Ar more removed (but see also Borra et al (2015)). We found 8Ar neurons display a rich set of response properties that were not frequently observed in FEF, suggesting an important distinction in how these two areas function during perception and action.

### Visual field representation

We found that 8Ar neurons in one hemisphere represent the entire visual field. This representation is achieved not only by an enhancement of activity above baseline (a more traditional receptive field), but also through spatially tuned suppression below baseline (an inversion of the traditional receptive field). These regions of excitation and suppression spanned both hemifields, such that a typical 8Ar neuron provided spatial information about the entire visual field rather than a small region typical of receptive fields in early visual cortex. Regions of excitation in 8Ar could be in either the contralateral or ipsilateral hemifield, with suppressive areas typically located directly opposite (by 180°). While nearby neurons in some cases had spatially similar receptive fields, we observed no distinct pattern of topographic organization across the electrode arrays (Figure 11). Taken together, these observations suggest a highly distributed representation of space in 8Ar.

**Figure 11:**
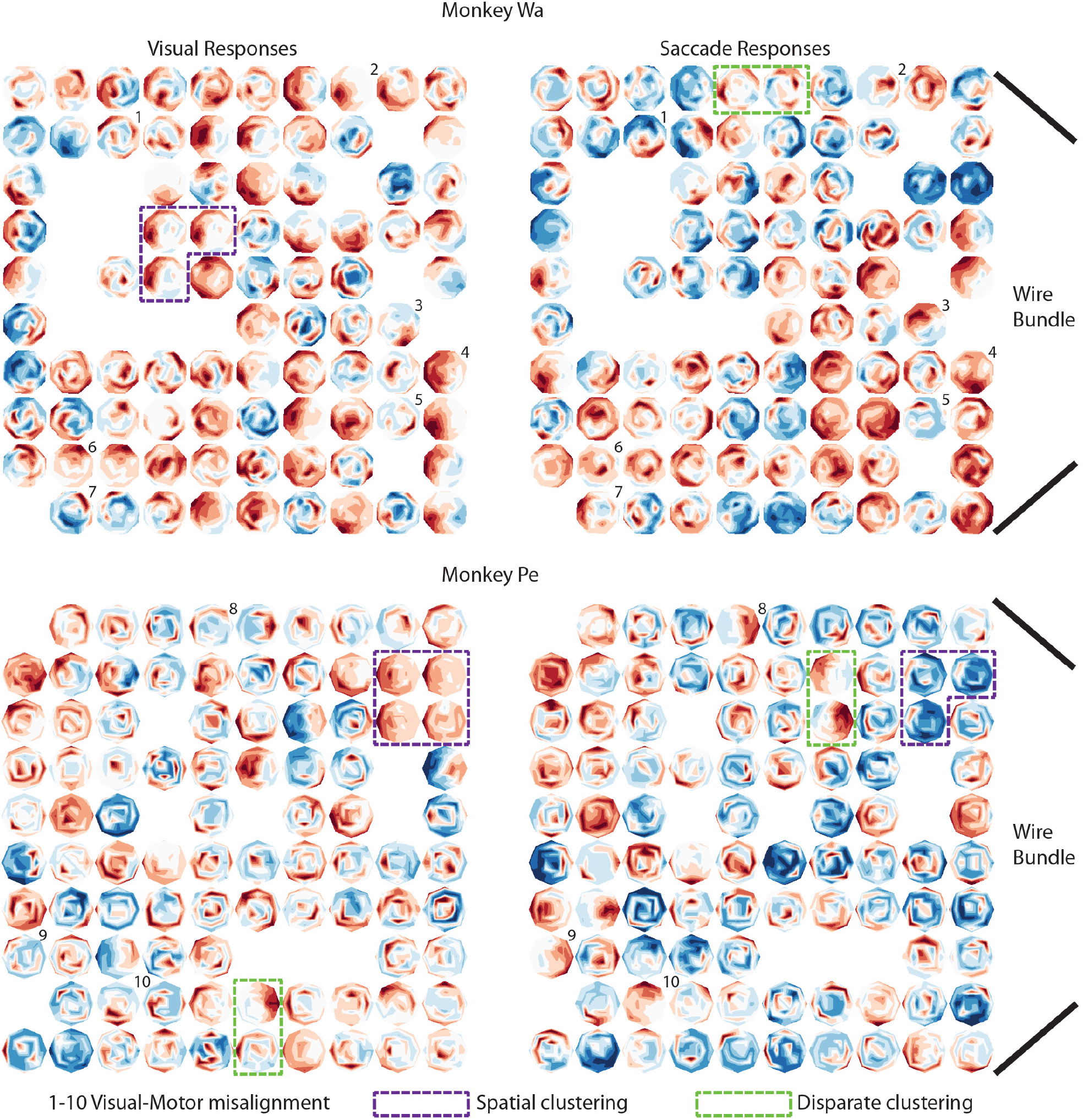
Topography of visual and motor responses in 8Ar. Response field maps for one example session in monkey Wa (top) and Pe (bottom) during the visual (left) and saccade (right) epochs. As with previous response field maps, red colors corresponded to activity above baseline, blue below baseline, and white near baseline. The spatial location of each neuron corresponds to its position on the electrode array. The arrays are oriented with the wire bundle coming from the right side of the figure (refer to Figure 1 for the orientation with respect to the brain). Very roughly, the bottom right of the arrays in these figures were most anterior, with the top right being medial. If multiple neurons happened to be recorded on the same electrode, the neuron with largest modulation depth (maximum firing rate – minimum firing rate) was used. Colored boxes highlight illustrative examples of neurons located physically near each other (i.e., recorded with adjacent electrodes) with the same tuning (purple) or disparate tuning (green). Black numbers identify neurons with mixed selectivity between the visual and motor epochs.

In visual and oculomotor areas, neurons primarily represent the contralateral hemifield. This includes FEF (Bruce & Goldberg 1985, Mayo et al 2015), LIP (Ben Hamed et al 2001, Blatt et al 1990, Patel et al 2010), SC (Cynader & Berman 1972, Goldberg & Wurtz 1972, Schiller & Koerner 1971), and SEF (Schlag & Schlag-Rey 1987). Some cortical areas show evidence of ipsilateral tuning, such as MT (Gattass & Gross 1981, Van Essen et al 1981), FST (Desimone & Ungerleider 1986), LIP (Dunn & Colby 2010), SEF (Schall 1991, Schlag & Schlag-Rey 1987), and IT (Ungerleider 1983), although ipsilateral responses in these areas are typically confined to regions of space that are just across the vertical meridian. Our findings of ipsilateral tuning are consistent with earlier reports in 8Ar of visual, delay, and saccadic responses (Funahashi et al 1989, Funahashi et al 1990, Funahashi et al 1991, Mikami et al 1982, Suzuki & Azuma 1983). Some previous studies in 8Ar have identified neurons that are suppressed below baseline, primarily opposite the receptive field (Bullock et al 2017, Kiani et al 2015), as well as a medial-lateral topography for visual eccentricity (Suzuki & Azuma 1983) that mirrors that in FEF (Bruce et al 1985). We did not observe a strong topography in our recordings, but did observe local clustering consistent with a previous report (Leavitt et al 2017), evident from our observation of nearby neurons with similar RFs (Figure 11, dashed purple outline). It is likely that some of the variability in previous observations of topography within 8Ar is due to sparse spatial mapping in concert with the rich spatial structure of excitation and suppression.

### Transition from visual to motor responses in 8Ar and FEF

Our comparison of 8Ar and FEF, from the presentation of a visual stimulus to the execution of an eye movement, revealed key differences in the properties of these two cortical regions that are highly interconnected (Huerta et al 1987, Stanton et al 1993, Stanton et al 1995). Visual latencies for 8Ar neurons were slower on average with a broader range than FEF, and the tuning across the population was weaker in all groups (visual, motor, visuomotor) compared to FEF. This led to lower overall decoding accuracy in 8Ar compared to FEF, due to the relatively poorer direction selectivity and reliability of 8Ar neurons. Importantly, we observed many 8Ar neurons changed their tuning between the visual and motor epochs, with a sometimes striking misalignment between their preferences in these two periods of time that was revealed by the dense spatial mapping protocol we employed. These results in 8Ar are consistent with a transition from visual to motor representations that are instantiated with a different mixture of neurons and activity. Evidence for a similar misalignment in preferences has been previously reported in both FEF (Sajad et al 2015, Sajad et al 2016) and SC (Sadeh et al 2015) neurons in head unrestrained monkeys, where visual and movement responses most strongly encode target and gaze, respectively. Our findings reflect an even more fundamental misalignment between visual and motor target signals in some neurons.

The alignment of visual and motor signals in a neuronal population could be beneficial in areas close to the motor output, as a direct mapping provides an efficient means for processing information and generating a rapid movement. This circuitry seems to be implemented in SC, where visual and movement activity is spatially aligned (Wurtz & Goldberg 1972). Given the strong, topographic descending projections from FEF to SC (Stanton et al 1988), and the observation that microstimulation in FEF elicits saccades at very low currents (Bruce et al 1985), such an alignment might also be expected in FEF. Indeed, when comparing FEF to 8Ar, visual and motor signals were more aligned in FEF. Why might different strategies be implemented in neighboring cortical regions that share involvement in important visuomotor behavior? Despite their proximity and interarea connectivity profile, lesions to FEF and 8Ar have resulted in differentiable deficits. In an anti-saccade task, for example, lesions in adjacent area 46 of PFC result in an increased percentage of errors (Pierrot-Deseilligny et al 1991, Ploner et al 2005) while in FEF there is an unchanged error rate but an increased saccade latency (Fukushima et al 1994). Another study which directly compared 8Ar and FEF during a distractor task, found the FEF code was more generalizable (in agreement with our findings), and that the 8Ar code morphs to account for the distractor and still preserve information about the stimulus (Parthasarathy et al 2017). A comparison of 8Ar with lateral intraparietal cortex (LIP), an oculomotor region strongly connected with FEF (Barbas & Mesulam 1981, Ferraina et al 2002, Medalla & Barbas 2006), found relatively stable decoding of LIP activity over time and a more dynamic 8Ar code that was more robust in the presence of distractors (Meyers et al 2018). Our observations of the tuning properties of these two regions, and their differing alignment between visual and motor codes, are consistent with 8Ar playing an important role in more flexible (and less reflexive) visuomotor behaviors. One possible advantage of misalignment between the code for a visual stimulus and for the execution of an eye movement could be to avoid one signal contaminating the other, enabling an animal to resist moving its eyes to the location of a salient visual stimulus.

### Interpretation of dynamic selectivity & implications for working memory models

The first studies investigating the neural correlates of working memory in PFC observed elevated spiking during the memory or delay period (persistent activity), and concluded this was the source of the working memory signal (Fuster & Alexander 1971, Kubota & Niki 1971). However, later work demonstrated many PFC neurons were transiently activated (Romo et al 1999, Warden & Miller 2007, Zaksas & Pasternak 2006) and at the population level, the code appeared to be not persistent, but dynamic (Barak et al 2010, Meyers et al 2008, Stokes et al 2013). This body of work has contributed to a vigorous debate on how working memory is represented in cortex, with some supporting a persistent model (Constantinidis et al 2018) and others a dynamic model (Lundqvist et al 2018). While both groups agree activity during the memory epoch is important and that there exists dynamic tuning at the level of single neurons, they differ in what aspects of the activity are proposed to underlie the working memory signal. One model suggests the working memory signal lies in a stable subspace, permitting a fixed population readout despite the dynamics of individual neurons (Murray et al 2017). Others suggest the dynamics are how the memory is encoded, either through an “activity silent” mechanism (Stokes 2015) or through sparse coordinated spiking facilitated by oscillations in local networks (Lundqvist et al 2016).

Our study remains agnostic to which is the appropriate model for working memory, and instead focuses on potential origins of dynamic delay activity in individual neurons and populations. We found that much of the dynamic evolution of activity in 8Ar could be explained by the transition between representations of perceptual input and motor output that involve different mixtures of neurons in the population. That is, the dynamics we observed were not random fluctuations in the activity of individual neurons during the delay period, but rather the transition between two separate spatial tuning functions for visual input and motor output. This suggests the 8Ar code is stable but dynamic, a similar interpretation to Spaak et al (2017). Because lateral PFC as a whole has been implicated in a wide range of sensory and cognitive behavior (Tanji & Hoshi 2008), this leads to the speculation that apparent dynamics in 8Ar may be explained by other task variables for which individual neurons are tuned, such as differing sensory input and motor output modalities, reward anticipation, spatial attention, and more. Dynamics at the level of individual neurons could be a natural consequence of the implementation of such a flexible input and output structure instantiated in an overlapping fashion in a population of neurons. In such an environment, stable population readouts might be achieved in a manner that allows the stored memory item to be separated from the other variables concurrently represented in the network (Murray et al 2017, Rigotti et al 2013).

### Limitations of this study

The design of the current study incorporated two choices that are worthy of discussion here. First, our targets were concentrated in the central 30° of visual angle (up to 15° saccade amplitudes in 8 directions) in our 8Ar recordings, and fixed at 10° saccade amplitudes (8 directions) in FEF (a compromise eccentricity effective in driving responses for many neurons in the region of our FEF electrode tracks). The seminal work measuring topography in 8Ar (Suzuki & Azuma 1983) reported a medial-lateral gradient with smaller, more foveal RFs located laterally and larger, more eccentric RFs located medially. In the region just dorsal to the principal sulcus, where our arrays were implanted, they recorded from some neurons with RFs that were centered beyond the extent of our target array. Thus, in some cases we may have recorded from 8Ar neurons in which we found weak or absent tuning merely because we did not present stimuli at the ideal location for each neuron. A hint of this can be seen in our response maps (Figure 11), where some neurons exhibit tuned responses at the edges of the tested region. Prior to establishing the target array tested here we did test each animal with larger eccentricity targets (up to 20-25°), and did not observe qualitatively better tuning at the population level, although some individual neurons did have tuning at those eccentricities. Moreover, our findings closely mirror recent studies of 8Ar (Bullock et al 2017, Kiani et al 2015), in terms of the implanted array locations and targets tested. Although the overall trend in this previous work supports a medial-lateral gradient in RF eccentricity, the relatively weak clustering of receptive field location is evident in the large scatter of RF centers we observed for even neighboring electrodes (Figure 11). Overall, our experiments were performed with the goal of identifying the response properties of neurons in 8Ar and FEF to a canonical set of stimuli for which the population was tuned, not necessarily the ideal stimuli for every neuron in the population. Thus, we cannot rule out that some of the differences we observed between 8Ar and FEF were due to the choices we made in the target locations we tested or sampling differences in our recordings of the two areas.

Our task was designed with a fixed 0.2 s pre-stimulus delay period prior to stimulus onset, and a fixed post-stimulus delay period of 0.5 s (or 0.6 s for FEF). We chose this fixed delay to maintain as constant a trial structure as possible, but this could have led to influence of the post-saccadic response on the baseline firing rate prior to the stimulus, as well as anticipatory saccade preparation toward the end of the delay period. With a much longer pre-stimulus delay (0.65 s), Bullock et al (2017) also reported suppressive regions in the RF often located opposite excitatory regions, and shifts in tuning between visual and perisaccadic epochs. Furthermore, we found suppressive regions often did not directly oppose excitatory regions (e.g., Figure 5A), making it unlikely that post-saccadic response alone could explain the suppression. Because our delay period was fixed, subjects could have begun saccade planning prior to the end of the delay period at fixation offset. This paradigm is known to contribute to early buildup in motor preparatory activity in the superior colliculus (Dorris & Munoz 1998), and thereby might have contributed to the rapid timescale over which we observed the transition from a visual to motor code in the neuronal population. However, such an effect would not have produced the pattern of excitatory and suppressive responses or the spatial shifts in RF tuning that we observed.

## Conclusions

Our results extend the literature in three key ways. First, due to the spatial resolution of our task, we were able to with high fidelity map the visual and saccadic responses of populations of 8Ar single neurons. We found a rich pattern of excitatory and suppressive responses in 8Ar that represented the entire visual field through contralateral and ipsilateral tuning. Second, the observed tuning shifted between epochs, quite often to the opposite hemifield, indicating what may be perceived as random dynamics are actually the result of a transition between a visual and motor code. Finally, we compared single neuron and population level responses from 8Ar and FEF, highlighting the unique dynamics of individual neurons and the population code in 8Ar even in a simple memory guided saccade paradigm. Taken together, these results demonstrate rich, but separate, visual and saccadic spatial representations in PFC underlie its flexible role in sensory and motor behavior.

## Acknowledgements

This work was supported by the National Institutes of Health (R01EY022928, R01MH118929, R01EB026953, P30EY008098); National Science Foundation (NCS 1734901); a career development grant and an unrestricted award from Research to Prevent Blindness; the Eye and Ear Foundation of Pittsburgh; and a National Institute of Health training grant (T32 EY017271-08 to S.B.K). We are grateful to Samantha Schmitt for technical assistance and Dr. Adam C. Snyder for his assistance with array implants and behavioral training for an unrelated experiment which facilitated this additional data collection in prefrontal cortex.

